# Gene expression identifies regional central nervous system vulnerability to heat

**DOI:** 10.64898/2026.06.22.733716

**Authors:** Susanna Pagni, Abderrezak Bouchama, Sanjay M Sisodiya

**Author notes:** Correspondence to: Sanjay M Sisodiya, Department of Clinical and Experimental Epilepsy UCL Queen Square Institute of Neurology Queen Square, Box 29, Queen Square, London WC1N 3BG, UK.

## Abstract

Heat-related illness is an increasing global health threat, with heatstroke representing its most severe form and frequently causing cerebellar injury characterised by selective Purkinje neuron loss. The molecular basis of this regional vulnerability remains unclear. Here, we integrated transcriptomic data from human heat exposure experiments, patients with heatstroke, primary human neurons, and cortical organoids to define conserved responses to heat stress and investigate determinants of cerebellar susceptibility.

Across datasets, heat stress induced a highly conserved transcriptional programme dominated by suppression of ribosome biogenesis, RNA processing, translation, and metabolic pathways, consistent with reduced biosynthetic and energy-demanding activity. Mapping these signatures to human brain atlases revealed that the cerebellum showed the lowest enrichment of downregulated genes, while heatstroke exhibited a distinct regional transcriptional pattern compared with experimental heat exposure. At the cellular level, granule and Purkinje neurons showed the strongest association with heat-responsive gene suppression. Purkinje-enriched genes were linked to synaptic organisation, neurotransmission, membrane excitability, and ion transport. Connectivity Map analysis identified compounds predicted to reverse the heatstroke transcriptional signature, including the mTOR inhibitor KU-0063794.

These findings identify a conserved molecular response to heat stress and suggest that selective Purkinje vulnerability reflects intrinsic metabolic and electrophysiological properties rather than preferential activation of canonical heat-shock pathways.

## Introduction

Heat-related illness encompasses a clinical spectrum of conditions arising when the body’s thermoregulatory capacity is overwhelmed by environmental heat, physical exertion, or both. Clinical manifestations range from relatively mild conditions such as heat cramps and oedema to life-threatening hyperthermia.^1^ At the severe end of this clinical spectrum lies heatstroke (HS), defined by a rapid elevation of core body temperature to above 40°C accompanied by central nervous system (CNS) dysfunction.^2^ Two major clinical phenotypes are recognised. Classic HS results from passive exposure to extreme environmental heat and typically occurs during heatwaves, especially in individuals with existing illnesses, whereas exertional HS develops during strenuous physical activity and predominantly affects otherwise healthy individuals.^2^

Between 2000 and 2019, heat-related mortality was estimated at approximately 489,000 deaths annually worldwide, with the greatest burden occurring in Asia and Europe.^3^ Major heatwave events have highlighted the scale of this threat. The European heatwave of 2003 resulted in more than 70,000 excess deaths and prompted widespread implementation of heat adaptation strategies across the continent.^4^ Nevertheless, the record-breaking European summer of 2022 was still associated with an estimated 61,672 heat-related deaths across 35 countries.^4^ The global burden of HS is also increasing rapidly. A systematic review of HS during the Hajj pilgrimage in Mecca documented profound physiological derangement in one of the largest reported cohorts of classic HS (2,632 people across 44 studies), with severe impairment of consciousness in more than half of affected individuals.^5^ HS incidence during Hajj has been reported to vary markedly according to climatic conditions, reaching a mean of 39.6 cases per 100,000 pilgrims during hot-cycle years compared with 2.1 per 100,000 during cooler periods.^6^ By comparison, population-based estimates from the United States suggest an annual heat-stroke incidence of approximately 1.34 emergency department visits per 100,000 population.^7^

Historically, HS was considered a relatively circumscribed hazard affecting specific high-risk populations, including military personnel, pilgrims exposed to desert climates, and elderly individuals during prolonged summer heatwaves. This epidemiological pattern is now changing. Climate change is increasing the frequency, intensity, and seasonal duration of extreme heat events worldwide, exposing populations with limited historical adaptation to severe environmental heat stress.^8^ Consequently, non-exertional HS is emerging as an increasingly important cause of morbidity and mortality in previously healthy people, even in temperate regions previously considered comparatively low risk.

The consequences of HS extend well beyond the acute event. Approximately one-third of survivors develop persistent neurological or peripheral tissue dysfunction, and the proportion of survivors unable to work or requiring institutional care increases markedly during long-term follow-up.^9^ Cardiovascular morbidity is also increased, with elevated long-term risks of ischaemic heart disease and heart failure following recovery.^10^ As extreme heat events become more common, the cumulative burden of chronic disability attributable to HS is likely to rise substantially, underscoring the need to better understand the mechanisms underlying heat-induced organ injury, with a view to prevention.

Among affected systems, the CNS occupies a central role in both the acute presentation and long-term consequences of HS. Delirium, seizures, altered consciousness and coma are defining clinical features across both classic and exertional HS forms.^2^ Neuroimaging studies performed months to years after HS have demonstrated persistent injury involving multiple brain regions, including the cerebellum, hippocampus, thalamus, and midbrain.^11^ Importantly, however, heat-induced neural injury is regionally selective rather than diffuse.

The cerebellum appears particularly vulnerable to heat-related injury. Among survivors of environmental, or non-exertional, HS with persistent neurological impairment, cerebellar dysfunction occurs in approximately 71% of cases,^12^ presenting as ataxia, dysarthria, dysmetria, nystagmus, and gait instability. Neuropathological studies identify selective degeneration of cerebellar Purkinje cells as the hallmark of fatal HS.^13^ This finding has been consistently replicated in both post-mortem and experimental studies, which demonstrate profound Purkinje cell loss, cerebellar demyelination, and persistent motor deficits following HS exposure.^14,15^

The mechanistic basis of this selective cerebellar vulnerability to hyperthermia remains unresolved. Proposed explanations include the high intrinsic metabolic demands of cerebellar circuitry, the sensitivity of Purkinje neurons to energetic and oxidative stress, and regional differences in cerebrovascular architecture that may limit local heat dissipation.^2,16^ However, whether the baseline transcriptional architecture of cerebellar tissue aligns with heat-responsive programmes to heighten cerebellar vulnerability remains unknown.

Human transcriptomic studies of blood provide an underlying rationale for this question. Controlled passive heat exposure in healthy volunteers induces a rapid cellular stress programme of heat-shock activation, repression of growth- and translation-associated pathways, mitochondrial dysfunction, and suppression of protein synthesis.^17^ In clinical heatstroke, this response is amplified into a disease-associated signature marked by broad chaperome activation, unfolded protein response, impaired oxidative phosphorylation and ATP production, and incomplete restoration after cooling,^18^ suggesting that injury may reflect failure of proteostatic and energetic adaptation when stress exceeds compensatory capacity rather than hyperthermia alone. Whether heatstroke-related amplification reflects a scaling-up of the same transcriptional architecture engaged during passive heat exposure, or qualitatively distinct molecular involvement of vulnerable structures, is unresolved.

To address this question, we integrated heat-driven transcriptional signatures derived from heat exposure studies of humans, primary human neurons, and cortical brain organoid models with spatially-resolved gene expression data from the adult human brain. Using this approach, we tested whether cerebellar regions exhibit preferential enrichment of heat-responding gene programmes relative to other CNS structures, thereby providing a potential molecular framework for their established clinical and neuropathological vulnerability to heat-related injury.

## Materials and methods

### Data sources

Peripheral blood mononuclear cell (PBMC) transcriptomic data were obtained from a controlled human heat exposure study (GEO Accession: GSE90763).^17^ In this dataset (“passive heat-stress PBMC”), samples were collected from 15 healthy adult volunteers at baseline, following 15 minutes of sauna exposure (75–78°C), and after a one-hour recovery period. To complement this experimental heat-exposure model with clinical heat illness, PBMC transcriptomic data from heatstroke patients were obtained from GEO Accession: GSE317880.^18^ This dataset (“clinical heatstroke”) comprised 19 patients presenting with heatstroke and 19 non-heatstroke controls recruited from the same hot environment; patient blood was sampled on arrival at the cooling unit before treatment and again after cooling.

Transcriptomic data from human primary neurons exposed to heat stress were obtained from GEO Accession: GSE132447.^19^ Neurons were subjected to heat shock at 45°C for 30 minutes, followed by recovery at 37°C. Differential expression outputs were used for downstream analysis, as provided by the original authors. RNA-sequencing data from cortical brain organoids were obtained from GEO accession GSE149797.^20^ Organoids (*n*=3 control, *n*=3 heat-shocked) were exposed to 41.5°C for 420 minutes.

### GTEx v11 human brain atlas

Tissue-level (ie bulk) transcriptomic data were obtained from the GTEx Analysis Release (2025-08-22, v11) (https://gtexportal.org/home/). Median transcript per million (TPM) values for protein-coding genes were extracted for 13 anatomically defined brain regions: amygdala, anterior cingulate cortex, caudate, cerebellar hemisphere, cerebellum, cortex, frontal cortex, hippocampus, hypothalamus, nucleus accumbens, putamen, spinal cord (cervical C-1), and substantia nigra.

### Differential expression analysis

Passive heat-stress PBMC microarray data were obtained as normalised log-transformed expression values. Probe identifiers were mapped to gene symbols using Bioconductor annotations, retaining the highest-variance probe for genes represented by multiple probes. Differential expression analysis was performed using limma with paired linear modelling to account for repeated sampling within individuals.^21^ Heat exposure recovery conditions (15 minutes and one hour) were compared with baseline. Benjamini–Hochberg (BH)-adjusted false discovery rate (FDR) <0.05 was considered significant.

Heatstroke patient microarray data were obtained as normalised log_2_-transformed expression values. Differential expression analysis was performed using limma, comparing heatstroke patients at arrival (pre-cooling) with non-heatstroke controls. BH-adjusted FDR <0.05 was considered significant.

Primary neuron data were obtained as processed differential expression outputs comparing untreated neurons with 1h, 3h, and 24h post-heat-shock timepoints, as provided by the original authors. Significance was defined at FDR < 0.05.

Cortical brain organoid RNA-sequencing data were analysed from TPM-normalised expression matrices comprising control and heat-shocked organoids. Expression values were transformed as log(TPM+1), and genes with low expression (mean log expression ≤1 in both conditions) were excluded prior to analysis. Differential expression analysis was performed using limma^21^. Genes with BH-adjusted FDR<0.05 were considered differentially expressed.

### Functional enrichment analysis

Functional enrichment analysis was performed using the clusterProfiler R package to identify over-represented Gene Ontology (GO) Biological Process (BP) terms. Upregulated and downregulated gene sets were analysed separately for each dataset. P-values were adjusted using the BH procedure. The top 15 enriched terms per condition were visualised using dot plots.

### Brain regional expression of heat-responding genes

Heat-responding DEGs were mapped to adult human brain gene expression using the GTEx v11 atlas across 13 brain regions. Expression values were log(TPM+1)-transformed, and genes with mean log(TPM+1) > 0.5 across tissues were retained (20,071 genes).

Because brain regions differ in baseline transcriptomic expression levels, with the cerebellum showing globally elevated expression, absolute expression values were not compared directly. Instead, we quantified where heat-responding genes ranked within each region’s own transcriptome. For each DEG set and direction of effect, within-region enrichment was defined as the probability that a randomly selected DEG was expressed at a higher level than a randomly selected background (non-DEG) gene, equivalent to the area under the receiver operating characteristic curve (AUC). AUC values were derived from the Mann–Whitney (Wilcoxon rank-sum) statistic (*W*) as AUC=W/(n_DEG×n_background)^22^. An AUC>0.5 indicates that the DEG set is expressed above the region-specific transcriptome-wide background, whereas AUC<0.5 indicates relative underexpression. Because the metric is rank-based within each region, it is insensitive to differences in baseline expression magnitude across regions and robust to extreme outlier genes. Within each DEG set and direction, regions were compared pairwise using BH-adjusted paired Wilcoxon signed-rank tests.

### Cerebellar cell-type expression of heat-responding genes

Cell-type-specific expression of heat-responding genes within the cerebellum was assessed using a published single-nucleus RNA-seq dataset of adult human cerebellum (GEO: GSE97930), comprising 5,601 nuclei from four neurologically normal individuals.^23^ Donors were aged 35–49 years (two female, two male) with post-mortem intervals of 13–24 h; none had a reported history of neurological disease or heat-related illness. The dataset included 11 transcriptionally-defined clusters, including two Purkinje populations (Purk1, Purk2), which were averaged within each donor to yield a single Purkinje value. Cell-type nomenclature was retained from the original publication; the “cerebellar” prefix for certain cell clusters reflects transcriptionally distinct populations identified by the authors through multimodal integration across brain regions; clusters lacking this prefix were not cerebellum-specific and showed expression profiles more closely aligned with corresponding non-cerebellar cell populations. For each donor–cell-type combination, unique molecular identifier (UMI) counts were summed across nuclei to form pseudobulk profiles, normalised to counts per million (CPM), and log(CPM+1)-transformed. Genes with mean log(CPM+1)>0.5 were kept as the background universe (15,148 genes). Within-cell-type enrichment was quantified using the same rank-based AUC metric as the regional analysis, providing intrinsic baseline correction for differences in overall expression and capture efficiency across cell types. For each DEG set and direction, we fitted a linear mixed-effects model, with donor as a random intercept to account for non-independence of cell types from the same donor. The significance of the cell-type fixed effect was assessed by likelihood-ratio tests, followed by pairwise cell-type comparisons using estimated marginal means with BH correction (FDR < 0.05).

### Granule–Purkinje differences in baseline expression of heat-responsive genes

To identify heat-responding genes that distinguish granule from Purkinje neurons, we used the GSE97930 cerebellar pseudobulk profiles described above. Baseline differential expression between granule and Purkinje neurons was computed across all expressed genes (mean log(CPM+1) >0.5; 19,230 genes) using limma with donor as a blocking factor. Heat-responsive DEGs (BH FDR<0.05) were then overlapped with granule–Purkinje DEGs (BH FDR<0.05), yielding the subset of heat-responding genes that also differ in baseline expression between the two cell types. For each gene in this overlapping set (heat-responding DEGs and granule–Purkinje DEGs), mean expression (log(CPM+1)) was calculated across the ten cerebellar cell types. Cell-type specificity was quantified using the Tau index,^24^ which ranges from 0 (uniform expression) to 1 (expression restricted to a single cell type). Genes differentially expressed between granule and Purkinje neurons with Tau ≥0.80 were considered cell-type-specific markers and excluded prior to enrichment analysis.^25^ The remaining DEGs were separated into granule-higher and Purkinje-higher subsets and tested for GO Biological Process over-representation using clusterProfiler (BH FDR < 0.05). Analyses were performed independently for each heat-responding dataset.

### Drug-repurposing connectivity analysis

To identify compounds capable of reversing the heatstroke transcriptional signature, drug–disease connectivity was assessed using the Connectivity Map/LINCS L1000 framework,^26,27^ via the Bioconductor signatureSearch package.^28^ The heatstroke signature was derived from the GSE317880 differential expression analysis described above. The query comprised the top 150 upregulated and top 150 downregulated genes (BH FDR<0.05, ranked by significance), following Connectivity Map conventions.^26^ Connectivity was assessed against the full LINCS L1000 reference dataset (8,140 compounds across multiple cell lines). Weighted connectivity scores (WTCS) were aggregated across cell lines and perturbation replicates to generate a normalised connectivity score (NCS), which was converted to a tau (_τ_) metric (range −100 to 100) relative to the reference distribution.^27^ Negative _τ_ values indicate transcriptional reversal of the query signature, with _τ_≤−90 considered a high-confidence reversal signature.^27^ Compounds profiled in fewer than two cell lines were excluded. Mechanism-of-action annotations were obtained from the Broad Drug Repurposing Hub^29^ and LINCS metadata.

The pipeline was benchmarked using a curated positive-control signature representing constitutive mTORC1 activation in tuberous sclerosis complex (TSC). The signature comprised the MSigDB Hallmark mTORC1 signalling gene set (upregulated) and the union of KEGG autophagy and lysosome pathways (downregulated), retrieved via Enrichr, with overlapping genes removed to ensure disjoint sets. This configuration reflects transcriptional consequences of TSC1/TSC2 loss and was expected to show strong negative connectivity to mTOR inhibitors, providing validation of reversal detection within the framework.

### Drug-repurposing connectivity analysis of granule–Purkinje heat-response differences

To explore whether any compound could shift the vulnerable Purkinje transcriptome towards the more resilient granule-cell state, the connectivity pipeline described above was applied to a granule–Purkinje-resolved signature. We initially sought to derive this signature from the heatstroke dataset alone, mirroring the primary connectivity analysis. Although this yielded a valid query, only 69 Purkinje-enriched genes remained after removal of cell-type-identity genes (Tau ≥ 0.80), resulting in a small and asymmetric signature relative to the conventional Connectivity Map query size of up to 150 genes per direction^28^ used in the primary analysis. We therefore defined the signature using genes that met both criteria: differential expression between granule and Purkinje neurons and heat responsiveness in at least two of the four independent heat-stress datasets. This yielded a conventionally sized query while ensuring that the signature reflected reproducible transcriptional differences. Genes expressed at higher levels in Purkinje neurons formed the query up-set (142 genes), whereas genes expressed at higher levels in granule neurons formed the query down-set (150 genes). Under this framework, strongly negative connectivity scores (τ ≤ −90) identify compounds predicted to oppose the Purkinje-associated expression pattern and shift transcriptional profiles towards the granule-associated state. Compounds profiled in fewer than two cell lines were excluded.

## Results

### Transcriptional responses to heat stress across human cellular systems

To characterise the transcriptional response to heat stress, we compared differential expression across the four heat-exposure datasets described above. In PBMCs from healthy volunteers undergoing a controlled sauna protocol, the present reanalysis detected no genes reaching the prespecified FDR threshold at the 15-minute timepoint. A broad, directionally balanced response (51% downregulated) emerged at one hour of recovery; this timepoint was therefore used as the sauna heat-response signature in all subsequent analyses. DEG counts differed from those reported in the original study, likely reflecting differences in normalisation and differential-expression methodology between the two analysis pipelines.^17^

PBMCs from heatstroke patients sampled on arrival (pre-cooling) relative to non-heatstroke controls from the same environment showed a larger response dominated by downregulated genes. In both neural systems, the transcriptional response was predominantly suppressive: primary cortical neuronal transcriptomes responded most extensively at one hour of recovery and then resolved over time, with the directional balance becoming more equal by 24 hours, whereas cortical brain organoids exhibited the largest and most strongly downregulated response of any system (Figure 1A).

**Figure 1:**
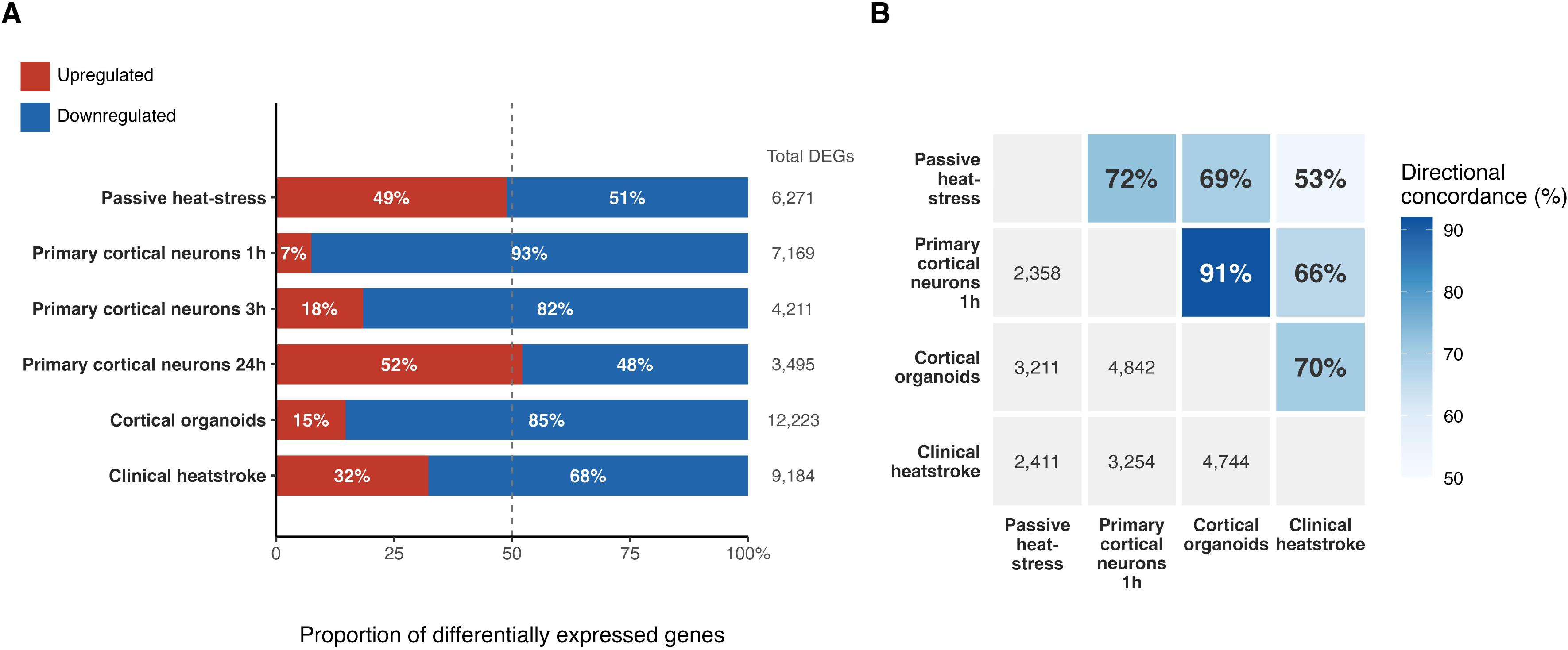
Directional asymmetry of heat-responsive transcriptional signatures across human cellular systems. (**A**) Stacked bars show the proportion of upregulated (red) and downregulated (blue) differentially expressed genes (DEGs) in each dataset, with total DEG count shown on the right. (**B**) Heatmap showing pairwise overlap of shared DEGs between datasets. The lower triangle shows the number of shared genes; the upper triangle shows directional concordance.

Pairwise comparison of the four datasets showed substantial shared DEG membership but variable directionality (Figure 1B). Concordance was highest between primary cortical neurons at 1 hour and cortical organoids, intermediate among the remaining experimental-system pairings, and consistently lowest for the clinical heatstroke cohort, which shared large numbers of DEGs with the other systems but showed only moderate directional agreement. Across all four datasets, 790 genes were differentially expressed in common, of which 436 were concordantly downregulated.

### Cross-dataset pathway overlap defines a conserved heat-stress transcriptional signature

Gene Ontology (GO) biological process enrichment analysis was performed separately for upregulated and downregulated DEG sets in each dataset. Among upregulated gene sets, enrichment yielded 287 significant GO terms in passive heat-stress PBMCs, 240 in clinical heatstroke PBMCs, 583 in primary neurons, and 320 in cortical organoids. With the exception of the heatstroke cohort, larger numbers of enriched terms were identified among downregulated genes, comprising 656 terms in passive heat-stress PBMCs, 177 in heatstroke PBMCs, 1,171 in neurons, and 1,080 in organoids.

To identify biological pathways that were consistently altered across experimental systems, GO enrichment results were integrated across the four independent datasets. Because the neuronal dataset contained three recovery timepoints, a GO term was considered neuron-associated if it reached FDR<0.05 at any timepoint. Among upregulated gene sets, no GO term was shared across all four systems. Canonical heat-shock and protein-folding–associated terms, including response to unfolded protein, chaperone-mediated and de novo protein folding, response to heat were strongly and highly enriched in three of the four systems, occupying the ten most significant upregulated terms in cortical organoids, the top ∼6% in heatstroke PBMCs, and similar positions in primary neurons, but were not enriched in the sauna-exposed PBMC dataset, so that no heat-shock term reached significance across all four systems.

By contrast, downregulated gene sets showed marked convergence across systems, with 72 GO terms shared between all datasets (Figure 2). These conserved pathways were dominated by fundamental biosynthetic and energy-associated processes, including ribosome biogenesis, RNA processing and splicing, cytoplasmic and mitochondrial translation, oxidative phosphorylation, mitochondrial respiratory-chain assembly, purine nucleotide biosynthesis, and proteasome-mediated protein catabolism (Figure 2). The 72 shared terms were among the most significantly enriched in every system: in three of the four datasets the single most significantly downregulated term was itself one of the shared set (ribosome or ribonucleoprotein complex biogenesis in neurons and heatstroke; purine-nucleotide metabolism in sauna-exposed PBMCs), and across systems the shared terms fell within the top quartile of each ranked enrichment list. Together, these findings indicate that the most conserved transcriptional feature across heat-stress systems was broad suppression of biosynthetic, translational, mitochondrial, and energy-associated cellular functions.

**Figure 2:**
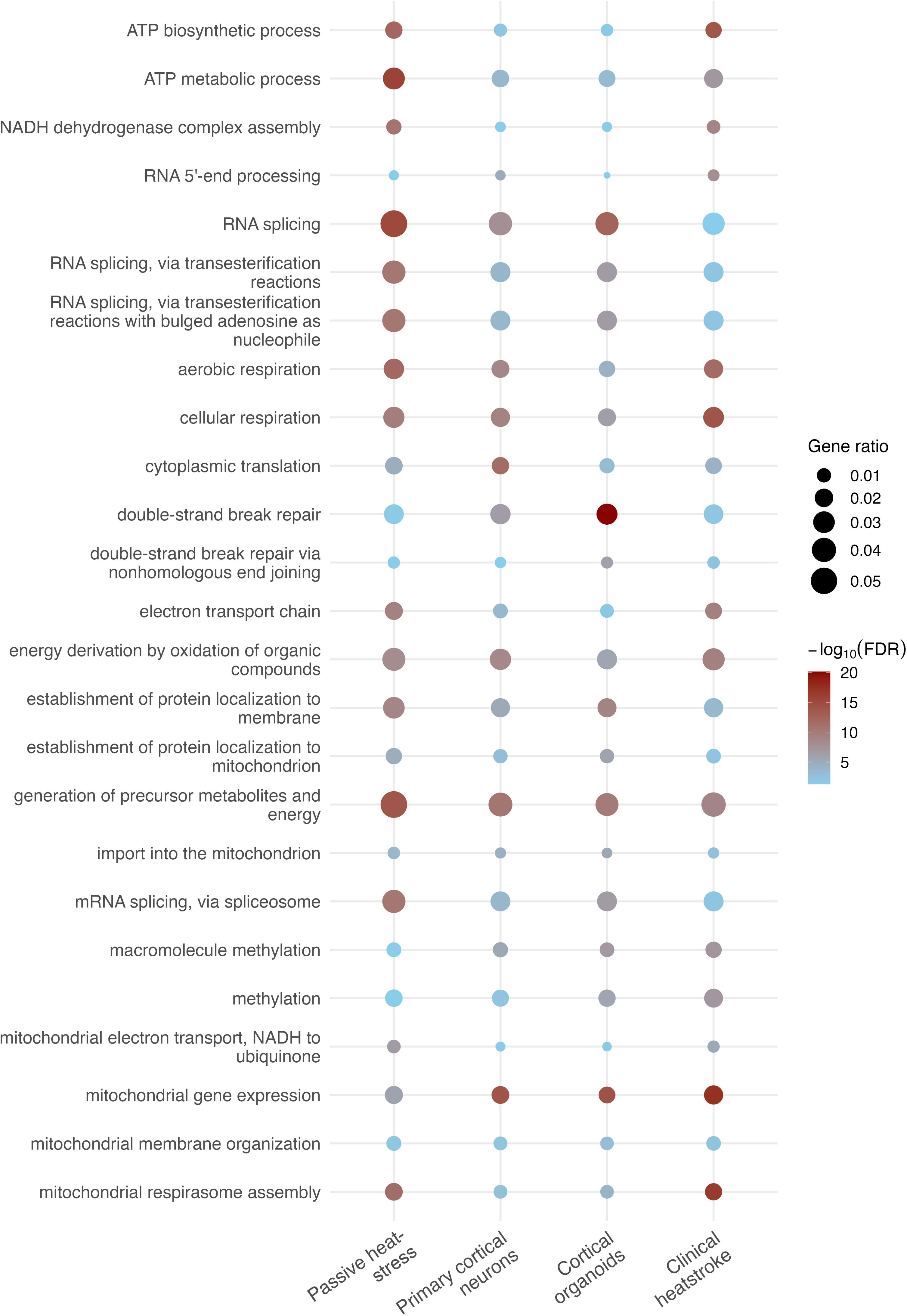
Conserved GO Biological Process terms across heat-stress experimental systems. Dot plot showing the GO Biological Process terms enriched among downregulated differentially expressed genes in the intersection of all four datasets. Dot size reflects gene ratio; colour represents the BH-adjusted P-value.

### Temporal pathway analysis identifies components of the neuronal heat-stress response

To dissect the temporal dynamics of transcriptional responses to acute heat shock in neurons, we performed time-resolved GO biological process enrichment analysis across the three post-heat-shock recovery timepoints (1h, 3h, 24h) in primary neurons.

#### Upregulated pathways: from acute stress defence to biosynthetic recovery

At one hour, the upregulated signature was dominated by canonical heat stress response terms, including response to heat, cellular response to heat, and response to temperature stimulus, indicating a strong early activation of heat shock pathways. This early response also co-activated stress-associated signalling pathways, including the p38 MAPK cascade, Wnt signalling, and membrane depolarisation. By three hours, heat shock response terms persisted, alongside emerging enrichment of hypoxia-related pathways, including response to hypoxia and response to decreased oxygen levels. p53-mediated signalling also appeared at this timepoint, consistent with activation of cellular stress and DNA damage surveillance pathways.

At 24 hours, heat shock terms remained present but were reduced in prominence, while biosynthetic and organelle-remodelling pathways became more prominent. These included ribonucleoprotein complex biogenesis, ribosome biogenesis, rRNA metabolic process, mRNA splicing via spliceosome, macroautophagy, mitochondrion organisation, and regulation of autophagy. The induction of ribosome biogenesis and translational machinery at 24 hours is notable given their relative suppression at one hour. Concomitant upregulation of autophagy-related pathways is consistent with activation of protein-quality-control and cellular-remodelling processes during recovery.

#### Downregulated pathways: from acute biosynthetic suppression to altered neuronal programmes

At one hour, the downregulated profile was dominated by biosynthetic and bioenergetic processes. Translational machinery was strongly suppressed, with enrichment of ribonucleoprotein complex biogenesis, ribosome biogenesis, rRNA metabolic process and processing, cytoplasmic and mitochondrial translation, and mitochondrial gene expression. Protein quality control pathways were also downregulated, including proteasome-mediated protein catabolism, protein polyubiquitination, and macroautophagy. Additional suppressed processes included vesicle trafficking, energy metabolism, endoplasmic reticulum proteostasis pathways, DNA replication, and RNA transport. By three hours, suppression of ribosomal and translational terms persisted but was attenuated, while downregulation of macroautophagy, DNA replication, and mitotic cell cycle phase transition continued. At 24 hours, ribonucleoprotein complex biogenesis, ribosome biogenesis, and cytoplasmic translation were no longer present among downregulated terms, and appeared in the upregulated signature, consistent with restoration of translational programmes (see above). In parallel, a distinct set of downregulated processes emerged, including synapse assembly, axonogenesis, regulation of neuron projection development, actin filament organisation, regulation of cell junction assembly, positive regulation of cell projection organisation, cognition, learning or memory, and glutamatergic synaptic transmission terms.

### Regional enrichment of heat-responding genes diverges between experimental systems and heatstroke

The regional enrichment pattern differed strikingly by direction of regulation, with the cerebellum occupying opposite extremes depending on the dataset (Figures 3, 4).

**Figure 3:**
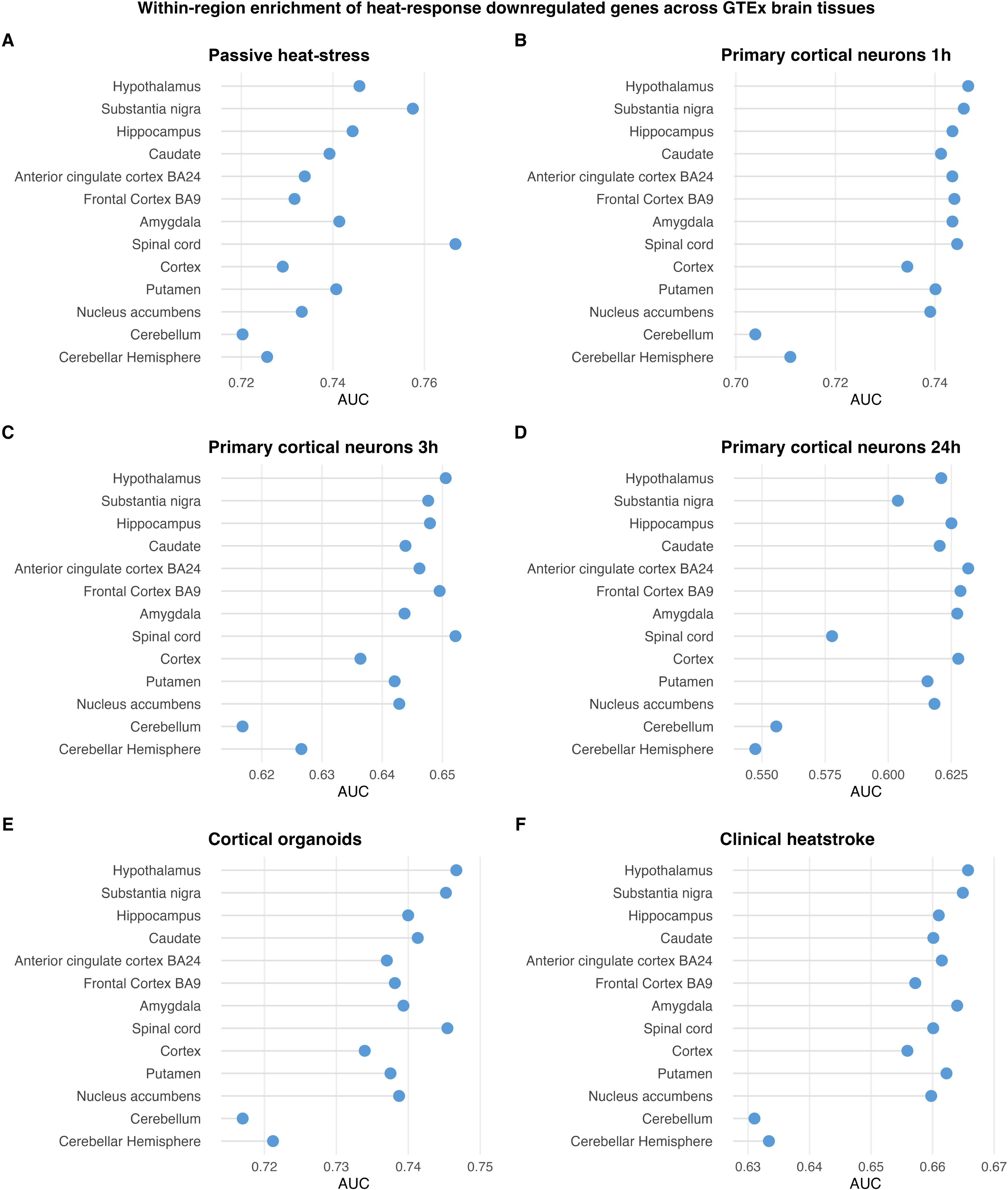
Within-region enrichment of downregulated heat-responding genes across human brain tissues. AUC scores for downregulated genes from each heat-stress dataset: (**A**) passive heat-stress PBMCs, (**B–D**) primary cortical neurons, (**E**) cortical organoids, (**F**) clinical heatstroke PBMCs, across brain regions. Each point represents one region. The dashed line at AUC = 0.5 indicates no enrichment.

**Figure 4:**
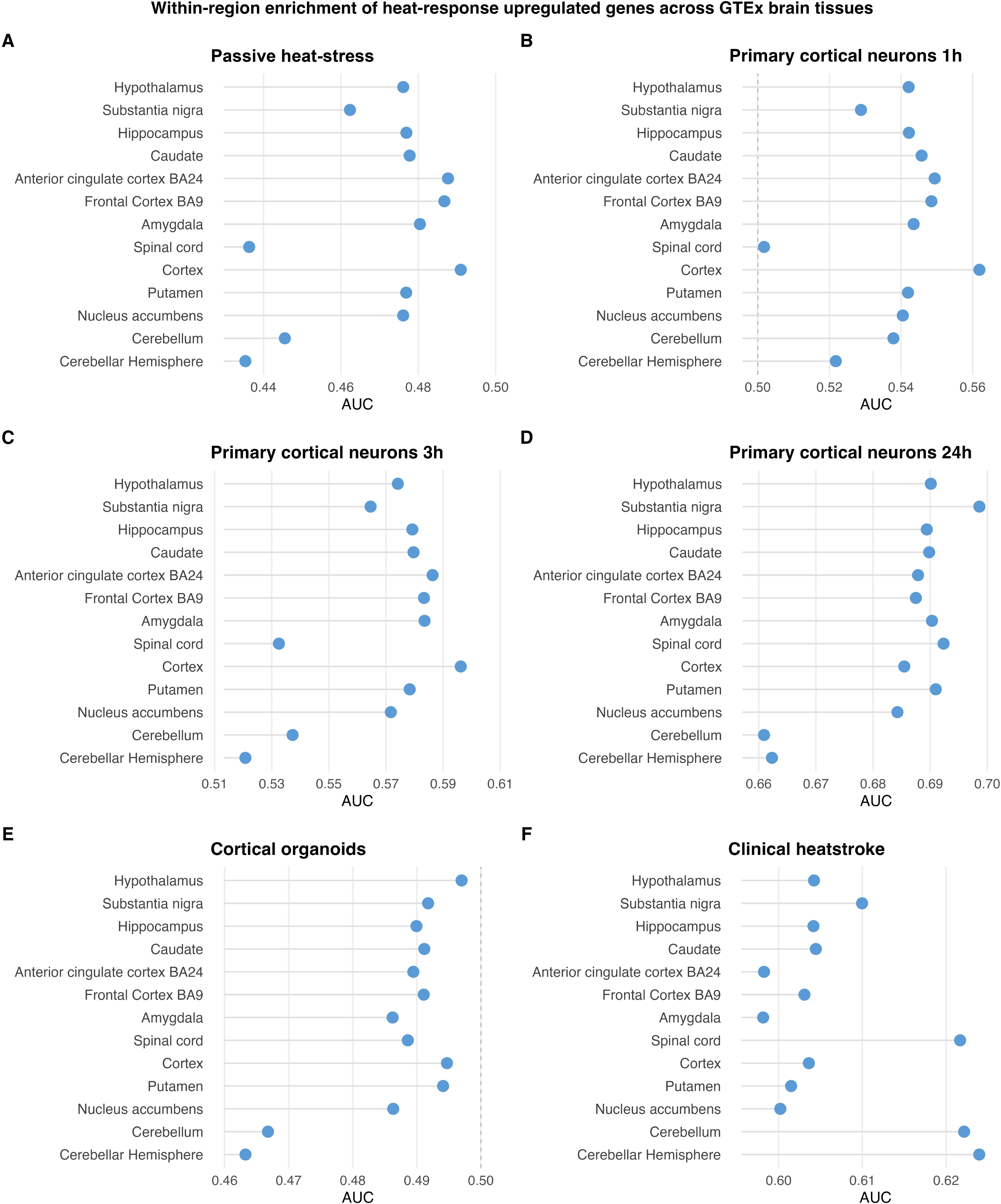
Within-region enrichment of upregulated heat-responding genes across human brain tissues. AUC scores for upregulated genes from each heat-stress dataset: (**A**) passive heat-stress PBMCs, (**B–D**) primary cortical neurons, (**E**) cortical organoids, (**F**) clinical heatstroke PBMCs, across brain regions. Each point represents one region. The dashed line at AUC = 0.5 indicates no enrichment.

For downregulated heat-response genes the pattern was uniform across all six datasets: the cerebellum ranked lowest of all brain regions in every dataset, with the cerebellar hemisphere consistently adjacent (Figure 3). In pairwise inter-region comparisons, the cerebellum ranked significantly below most of the other regions in every experimental dataset except the smallest DEG set (primary neurons, one-hour upregulated; *n* = 415), where it remained numerically lowest but did not reach significance.

For upregulated genes the pattern diverged by dataset. In the passive heat-stress PBMC, primary-neuron, and cortical organoid signatures, the cerebellum ranked at or near the bottom (ranks 10–13 of 13), with the highest enrichment observed in cortical regions, including cortex, anterior cingulate cortex, and frontal cortex. In the clinical heatstroke PBMC, however, this relationship was sharply inverted: the cerebellum and cerebellar hemisphere were the two highest-ranked regions of all analysed (Figure 4), significantly above the other regions in pairwise comparisons.

### Cerebellar cell-type expression of heat-responding genes largely follows a neuron–glia axis

For upregulated heat-responding genes, granule and Purkinje neurons ranked among the lowest-enriched cell types across the passive heat-stress PBMC, primary-neuron (one hour, three hours), and organoid signatures, while glial and vascular-associated cell types ranked highest, most consistently astrocytes. Two datasets departed from this pattern: primary neurons at 24 hours, where neurons ranked highest, reflecting the neuronal process dominance of that late-recovery gene set, and, importantly, the heatstroke cohort, where granule and Purkinje neurons were instead the two highest-enriched cell types (Figure 5). For downregulated genes the pattern was reversed and uniform across all six datasets including heatstroke: granule and Purkinje neurons showed the highest enrichment in every dataset (granule first, Purkinje second; heatstroke AUC 0.52 and 0.51, respectively), while glial cell types ranked lowest (Figure 6). Granule and Purkinje neurons tracked each other closely across all datasets and regulatory directions, with granule neurons consistently showing the more extreme effect. Pairwise comparisons confirmed that the strongest significant differences occurred along the neuron–glia axis.

**Figure 5:**
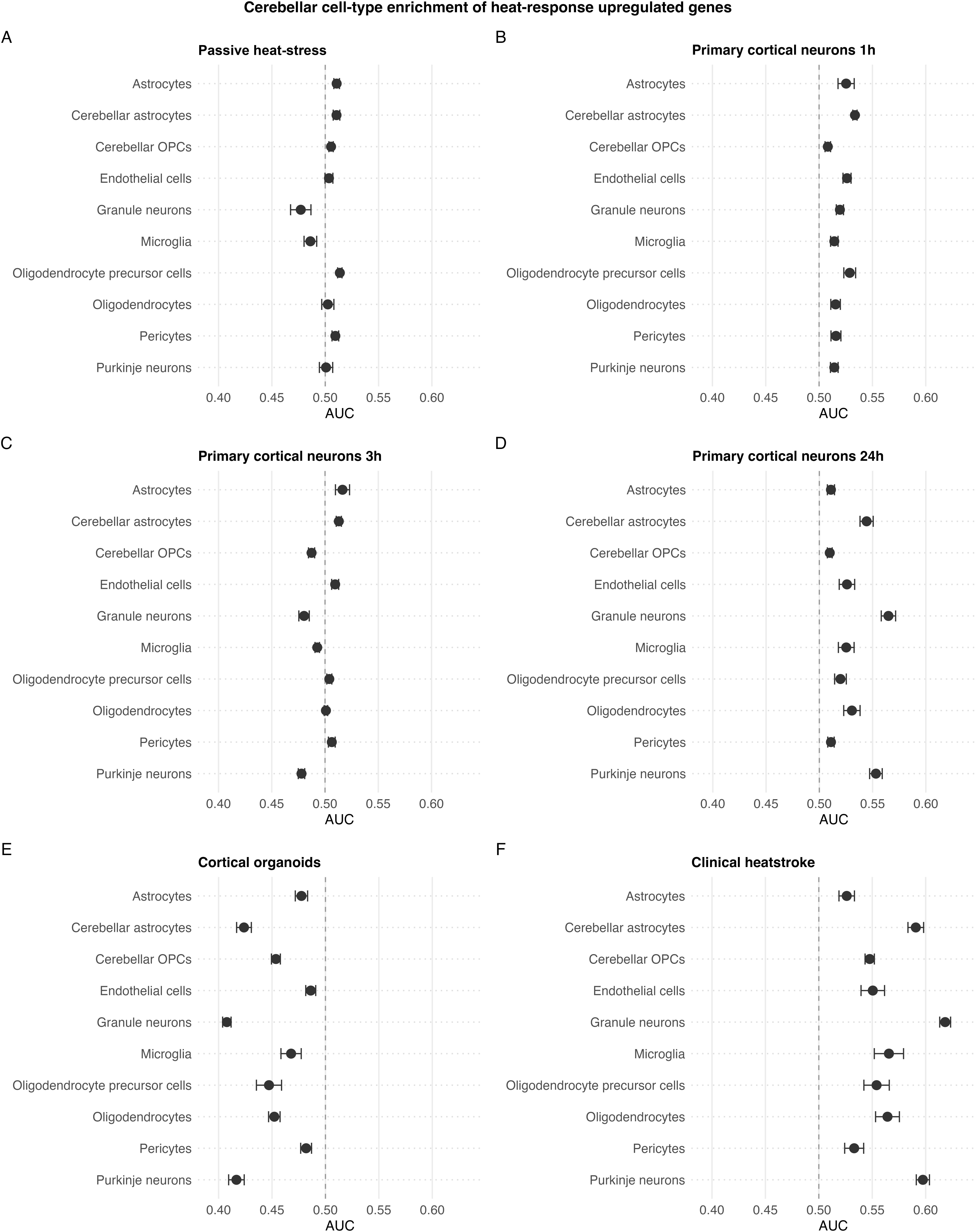
Within-cell-type enrichment of upregulated heat-responding genes across cerebellar cell types. Marginal mean AUC per cell type from the linear mixed model, with ± 1 SE bars, for upregulated genes from each heat-stress dataset: (**A**) passive heat-stress PBMCs, (**B–D**) primary cortical neurons, (**E**) cortical organoids, (**F**) clinical heatstroke PBMCs, across cerebellar cell types. The dashed line marks AUC = 0.5, which indicates no enrichment.

**Figure 6:**
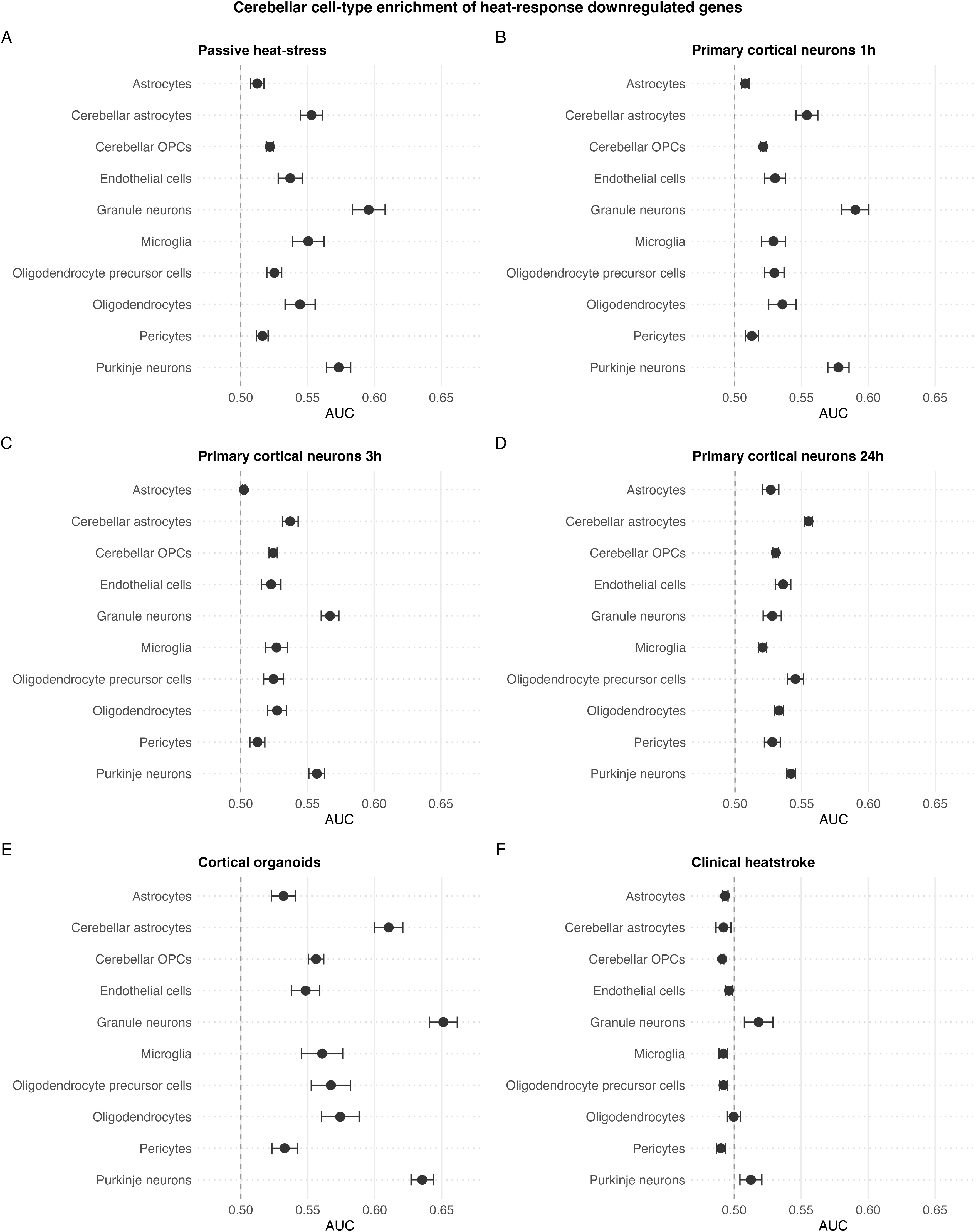
Within-cell-type enrichment of downregulated heat-responding genes across cerebellar cell types. Marginal mean AUC per cell type from the linear mixed model, with ± 1 SE bars, for downregulated genes from each heat-stress dataset: (**A**) passive heat-stress PBMCs, (**B–D**) primary cortical neurons, (**E**) cortical organoids, (**F**) clinical heatstroke PBMCs, across cerebellar cell types. The dashed line marks AUC = 0.5, which indicates no enrichment.

### Granule versus Purkinje neuron expression of heat-responding genes

Cerebellar heat injury is characterised by prominent Purkinje neuron loss with relative sparing of granule neurons.^30^ To investigate whether this differential vulnerability may be reflected in baseline transcriptional differences, we examined heat-responding genes that were also differentially expressed between granule and Purkinje neurons. After excluding cell-type identity genes, the Purkinje-higher DEG set showed substantial and reproducible functional enrichment across all four heat-response datasets, particularly for synaptic and excitability-related processes, including synapse organisation, chemical synaptic transmission, regulation of membrane potential, ion transmembrane transport, and cell junction assembly. In contrast, the granule-higher DEG set, despite containing approximately five- to sevenfold more genes than the Purkinje-higher set, showed little functional convergence, with only two enriched GO terms in the primary cortical neuron dataset (Figure 7).

**Figure 7:**
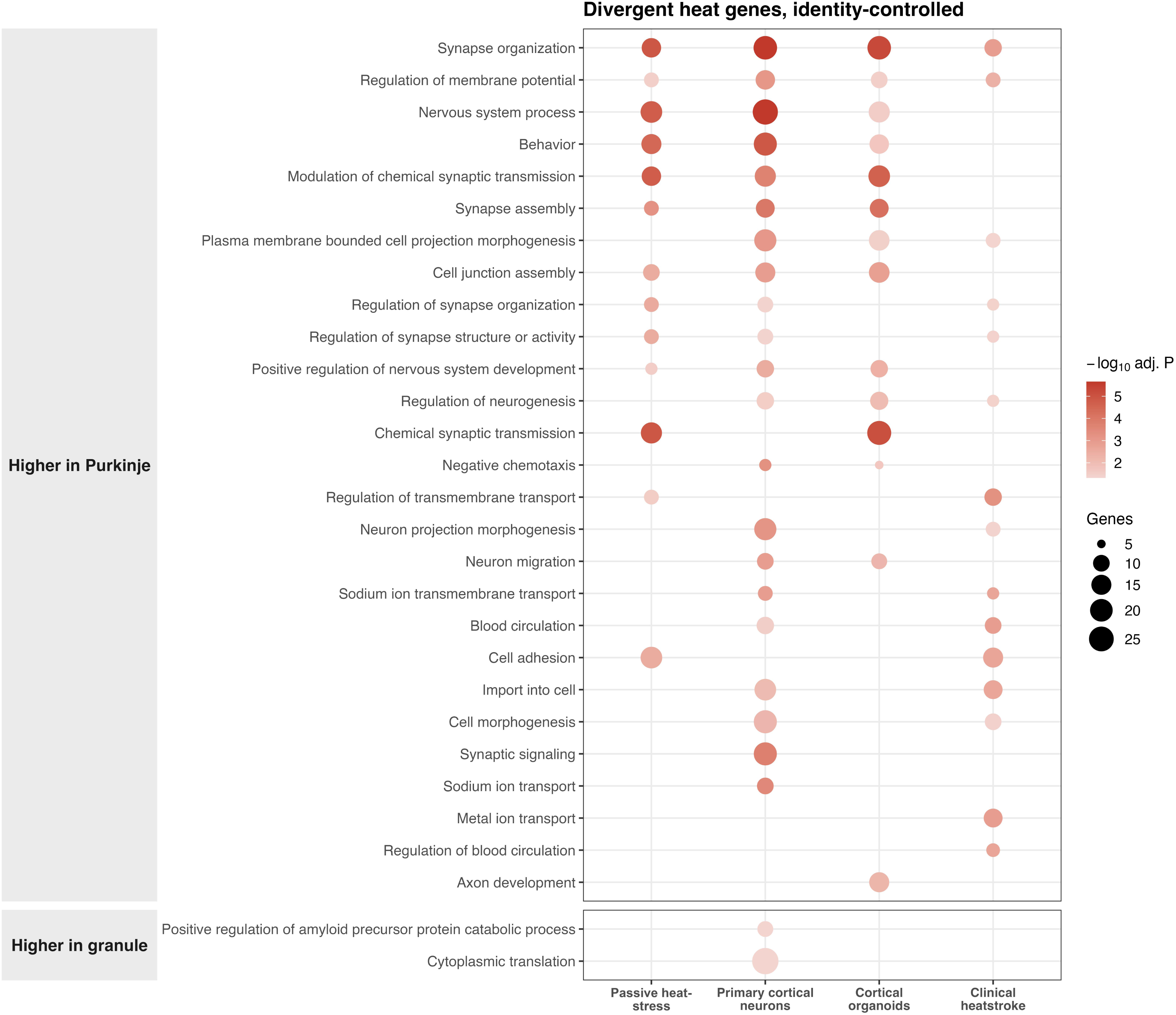
Heat-responding genes differentially expressed between granule and Purkinje neurons. GO BP enrichment analysis of heat-responding genes that were also differentially expressed between granule and Purkinje neurons after exclusion of cell-type identity genes. Dot size indicates the number of genes annotated to each term and colour indicates the BH-adjusted P-value.

### Drug-repurposing connectivity analysis of the heatstroke transcriptomic signature

Connectivity Map analysis identified 39 compounds that reversed the heatstroke transcriptional signature at the conventional significance threshold. However, most had been profiled in only a single LINCS cell line, precluding assessment of reproducibility across cellular contexts. Restricting the analysis to compounds tested in at least two cell lines reduced the set to eight candidate reversers. Among these, four compounds had annotated mechanisms of action: the mTOR inhibitor KU-0063794 (_τ_ = −92; eight cell lines), the SIRT1/2 inhibitor salermide, the acetylcholinesterase inhibitor edrophonium, and the cathepsin inhibitor SID-26681509. Several uncharacterised compounds showed even stronger negative connectivity scores, reaching _τ_=−98 (Figure 8); These correspond to structurally defined screened molecules without annotated targets or mechanisms, and are not currently tractable for in silico pharmacological interpretation or repurposing prioritisation. As a positive control, the pipeline correctly identified FDA-approved mTOR inhibitors as strong reversers of a tuberous sclerosis complex/mTORC1 activation signature, with temsirolimus and sirolimus achieving _τ_ scores of −98 and −94, respectively (Figure 8).

**Figure 8:**
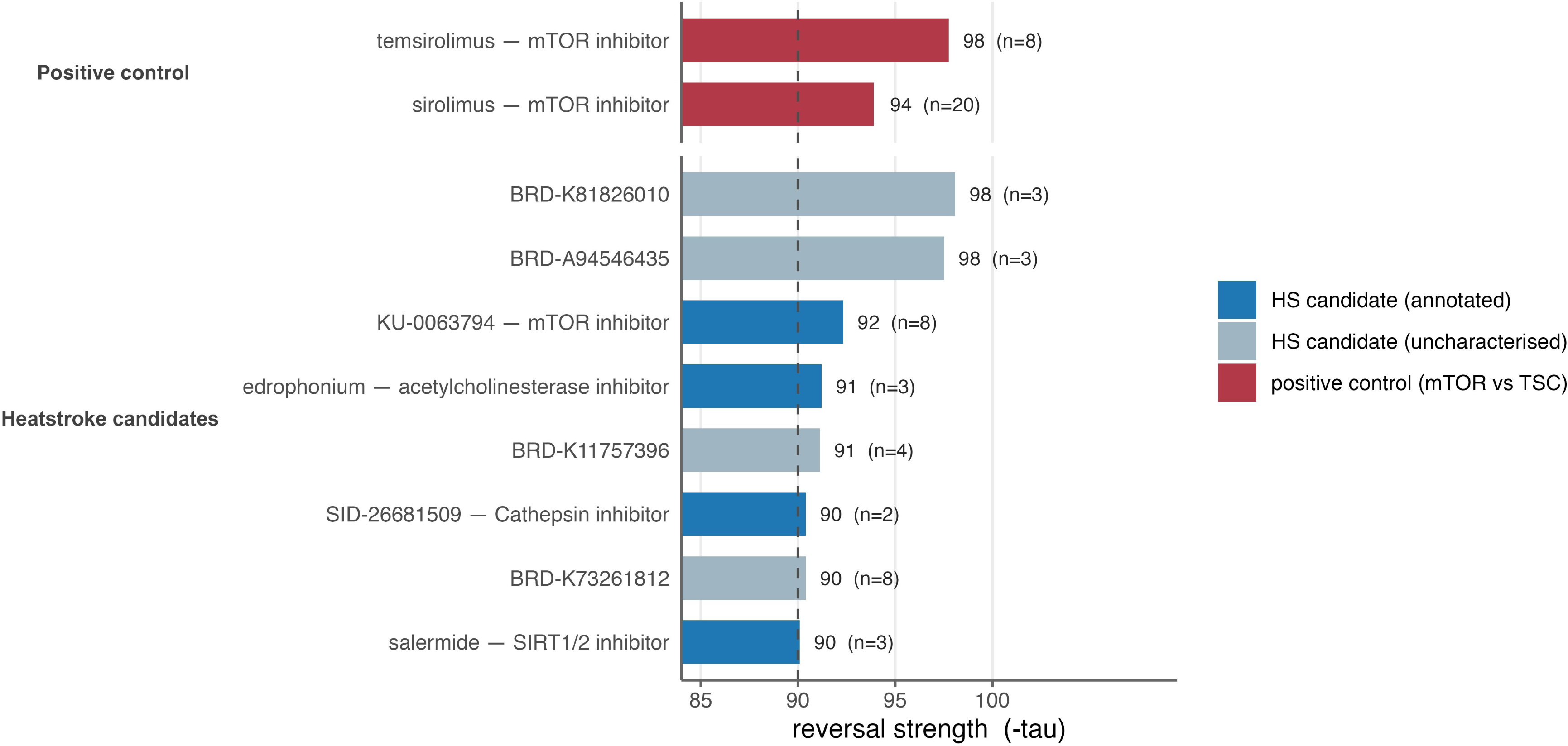
Connectivity Map analysis identifies compounds predicted to reverse the heatstroke transcriptional signature. Horizontal bars show Connectivity Map tau scores, where more negative values indicate stronger reversal of the heatstroke transcriptional signature. *n* denotes the number of independent LINCS cell lines in which each compound showed a reversing effect. The dashed line indicates the conventional Connectivity Map threshold for a strong inverse connection (_τ_=−90). FDA-approved mTOR inhibitors identified in the positive-control analysis of a tuberous sclerosis complex/mTORC1 activation signature are shown in red. Compounds predicted to reverse the heatstroke signature are coloured according to whether a mechanism of action is annotated in Drug Repurposing Hub/LINCS metadata.

### Connectivity mapping of granule–Purkinje heat-response differences

Applying the connectivity framework to the granule–Purkinje-resolved signature identified 17 compounds meeting the high-confidence reversal threshold (_τ_ ≤ −90), five of which were reproducible across multiple cell lines. Four were uncharacterised screening compounds lacking annotated molecular targets, while the single characterised candidate was trequinsin, a phosphodiesterase inhibitor that reproducibly reversed the signature across nine cell lines (_τ_ = −91).

## Discussion

The selective vulnerability of the cerebellum to heat injury is one of the most consistent features of heatstroke neuropathology,^13,14^ yet its molecular basis has remained poorly understood. A mechanistic understanding could help prevent acute and chronic sequelae of heatstroke, and potentially inform environmental adaptation measures.

This study integrates transcriptomic data from primary neurons, cortical organoids, human passive heat exposure and clinical heatstroke to investigate the molecular basis of heat stress and its relevance to selective cerebellar vulnerability. Across these systems, the analysis reproduced the archetypal heat-stress response, which combines induction of canonical stress-defence pathways with the expected predominance of transcriptional repression, as cells conserve and reallocate energy by suppressing growth, translation, and biosynthesis while prioritising protein protection and survival.^31,32^

Four principal findings emerged. First, the most conserved cross-system feature was broad suppression of biosynthetic and energy-associated processes. Second, there is a temporal programme of gene expression response in neurons. Third, heatstroke reproduces the core programme but displays a distinct regional and cellular architecture, particularly within the cerebellum: whereas cerebellar regions ranked low for upregulated heat-responsive genes in the experimental systems, they became the highest-ranked regions in clinical heatstroke. Fourth, extending this regional inversion to cerebellar cell types, the upregulated clinical heatstroke signature showed enrichment in cerebellar regions and cerebellar neuronal populations including granule and Purkinje neurons. The granule–Purkinje analysis suggested that selective Purkinje vulnerability was associated mainly with synaptic, excitability, and ion-handling programmes rather than canonical heat-shock defence pathways.

The most consistent feature of the heat-stress response was the convergence of downregulated pathways across all datasets. Despite substantial differences in cell type, experimental design, and clinical context, 72 GO terms were shared between all systems, predominantly involving ribosome biogenesis, RNA processing, translation, and related biosynthetic and metabolic functions. These shared pathways ranked among the most strongly downregulated processes in each system, indicating that the observed convergence reflects a conserved suppressive arm of the heat-stress response.^31,32^ In the context of our findings, the importance of this conserved programme is not that it redefines heat-stress response, but rather that it provides a common stress background against which cerebellar differences can be interpreted. Across systems, the magnitude of the response was particularly pronounced in cortical organoids, which showed the largest and most strongly downregulated transcriptional response of any system. This may reflect the fact that organoids recapitulate considerable cell-type and architectural complexity in vitro, yet develop in the absence of systemic support present in vivo, including vascular perfusion, immune interactions, and metabolic and thermoregulatory buffering. The combination of size and advanced intrinsic complexity with the absence of such extrinsic support may leave organoids particularly vulnerable to hypoxia and cell death and, potentially, thermal insult, amplifying the suppressive transcriptional response relative to the other systems.^33^

In contrast, the induced component of the heat-stress response did not reach a four-way intersection. Canonical chaperone and protein-folding-related pathways were among the strongest induced processes in the neuronal, organoid, and heatstroke datasets, yet were absent from the sauna-exposure PBMC dataset (one hour after sauna exposure, potentially after peak heat-shock induction). This heterogeneity could reflect differences in cell type, heat exposure, recovery stage, and the level of physiological organisation represented by each model. Whereas PBMCs were exposed to heat within an intact organism with complex thermoregulatory and systemic responses, neuronal cultures and organoids were studied in more reductionist experimental settings. These findings suggest that induced heat-stress programmes may be more context-dependent than the conserved suppressive response.

Beyond differences between systems, the neuronal time-course suggests that the heat-stress response unfolds in distinct phases. Early responses were characterised by canonical stress-response pathways, whereas later time points showed increasing suppression of neuronal and synaptic functions. This pattern suggests that restoration of basic cellular homeostasis may precede recovery of neuronal function. Such a dissociation may help explain why neurological injury can evolve despite apparent physiological stabilisation following acute heat stress, and is consistent with experimental and clinical observations of delayed neurological impairment after heatstroke.^14,15^ These findings also have potential therapeutic implications, suggesting that neuronal stress responses in heatstroke may extend beyond the period of initial physiological recovery and warrant further investigation in relation to the timing and targets of intervention.

An important question is whether heatstroke represents the severe end of a quantitative heat-stress continuum or a biologically distinct state. Our findings support elements of both models. Heatstroke clearly shares the conserved downregulated programme observed in experimental heat exposure, indicating preservation of the core cellular response to heat stress. However, the regional and cell-type analyses revealed a striking divergence in the organisation of the induced response. Across the experimental datasets, cerebellar regions consistently ranked among the lowest for enrichment of upregulated heat-responsive genes, and cerebellar neurons similarly showed relatively weak enrichment compared with glial populations. In the heatstroke cohort, this pattern was reversed: cerebellar regions became the most enriched tissues, and granule and Purkinje neurons the most enriched cell types. A purely quantitative model of heatstroke would predict amplification of the same transcriptional architecture observed during experimental heat exposure. Instead, the architecture itself changed. These findings suggest that heatstroke comprises the canonical heat-stress response together with additional pathological processes that preferentially engage cerebellar neuronal programmes. Heatstroke presents a complex acute and chronic picture, and our findings suggest that some of its distinct features may be underpinned by transcriptomic differences emerging only in extremis.

The cerebellar findings are of key relevance in light of the well-established, but mechanistically unexplained, neuropathology of heatstroke: selective cerebellar injury, especially involving Purkinje neurons, is a consistent neurological feature of heatstroke and can result in persistent ataxia despite survival from the acute illness.^14,30^ Overall, granule and Purkinje neurons showed broadly similar extents of heat-responding gene expression despite their marked contrasting vulnerability to heat. However, this similarity was observed at the level of aggregate transcriptional response rather than shared regulation of the same individual genes. We therefore compared these populations directly to identify transcriptional features that might underlie their differential vulnerability. After removal of highly cell-type-specific (canonical, cell-type identity) genes, no heat-shock, chaperone, or proteostasis-related pathways were preferentially associated with either population. Instead, genes expressed at higher levels in Purkinje neurons consistently converged on synaptic organisation, neurotransmission, membrane excitability, ion transport, and related processes. By contrast, the substantially larger granule-higher gene set showed little functional convergence across datasets. The absence of preferential chaperone pathway enrichment in Purkinje neurons should not be interpreted as a lack of proteostatic defence. Experimental hyperthermia studies in rabbit cerebellum showed that inducible hsp70 is rapidly upregulated in glia and granule neurons but not in Purkinje cells, which instead maintain high constitutive levels of hsc70 and exhibit only a blunted and delayed inducible response.^34^ Human post-mortem studies independently identified a similar pattern, with Hsp70-family transcripts prominent in granule neurons but absent from Purkinje neurons, consistent with an intrinsic cell-type difference rather than an agonal effect.^35^ Together, these findings support a model in which Purkinje neurons rely predominantly on constitutive proteostatic mechanisms, with a comparatively limited inducible reserve. Such a strategy may be sufficient to support their sustained physiological demands under normal conditions, yet prove inadequate when confronted with the additional proteotoxic and energetic burden imposed by extreme heat stress: Purkinje neurons can be viewed as high-demand neurons with high constitutive protection but limited inducible reserve.

A key limitation of the granule-versus-Purkinje transcriptomic comparison is that it relies on baseline transcriptional differences between granule and Purkinje neurons and therefore cannot capture genes that are similarly expressed at rest but diverge in their heat response. Such effects cannot be resolved from the present data, as the heat-response and granule-versus-Purkinje comparisons were derived from separate experimental systems, and heat stress was not applied to the two cell types in a matched, cell-type-resolved design. To assess dependence on this filter, a robustness analysis removing this constraint recovered a modest proteostasis and RNA-processing signal in the granule-higher set, concordant with the biosynthetic signal of granule neurons established by the baseline gene set enrichment analysis, and did not alter the Purkinje synaptic signal.

Purkinje neurons are among the largest and most metabolically-demanding neurons in the brain; they are characterised by extensive dendritic arbours, sustained firing activity, substantial calcium flux, and high energetic requirements.^36^ Quantitative estimates indicate that individual Purkinje neurons consume approximately 1.24×10¹ ATP molecules per second, placing them among the most energy-demanding cell types characterised in the human brain, and substantially exceeding cerebellar granule neurons (≈1.72×10 ATP molecules per second).^16^ Computational modelling further predicts that Purkinje cells consume ∼60-fold more energy than granule cells and nearly four times more than cortical pyramidal neurons.^37^ Similar properties have been proposed to underlie their selective vulnerability in several neurodegenerative and excitotoxic disorders.^38–40^ The present results suggest that heat stress may expose the same underlying liability. In this framework, heat stress disrupts cellular homeostasis in both granule and Purkinje neurons, but the greater functional demands imposed by synaptic transmission and membrane excitability render Purkinje neurons less able to tolerate that disruption.

Looking to protecting Purkinje cells, to help reduce both acute and chronic brain consequences of HS, the Connectivity Map analysis identified several compounds predicted to reverse the heatstroke transcriptional signature, including the mTOR inhibitor KU-0063794. mTOR signalling occupies a central position in the regulation of translation, ribosome biogenesis, metabolism, and autophagy, processes that emerged repeatedly in the conserved heat-stress programme.^41^ Although these results are hypothesis-generating and do not establish therapeutic efficacy, they highlight mTOR-related pathways as candidates for further investigation in experimental models of heat-related injury. The in-silico analyses presented here, though derived from laboratory and clinical experimental data ultimately cannot by themselves securely distinguish, for example, true resilience or vulnerability pathways, or indeed epiphenomena, but point to possible mechanisms worthy of further exploration. Likewise, the data cannot securely determine which drug treatments might boost Purkinje cell protection. Noting this caveat, we did undertake a comparison of presumed non-cell-identity differentially expressed genes differing between granule and Purkinje neuronal types and drugs that might potentially reverse the unique, presumably deleterious Purkinje neuronal vulnerability signature. In this context, it is interesting to note that the single identified named compound, trequinsin, has already been studied as a potential neuroprotective agent.^42^

Several limitations of this work should be acknowledged. The regional and cell-type analyses were performed using baseline transcriptomic data from neurologically normal adult human tissue rather than heat-exposed brain tissue. Consequently, these analyses identify intrinsic transcriptional features that may influence susceptibility to heat-related injury rather than directly measuring cellular responses to heat stress in vivo. None of the experimental datasets used to derive heat-responsive gene sets originated from cerebellar tissue: the PBMC datasets were generated from peripheral blood, while the primary neuron and organoid datasets were derived from cortical regions. Although this limits direct tissue matching, the cerebellar signal was nevertheless detected using non-cerebellar datasets, arguing against a simple tissue-specific artefact. Inclusion of cortical brain regions in the atlas analysis also enabled direct comparison with areas that are relatively spared clinically. Furthermore, the heat-response signatures were derived from multiple experimental systems that differ in cellular composition, heat-exposure, and sampling times. While this heterogeneity may reduce sensitivity to system-specific effects, it also strengthens confidence in biological processes that reproduce across datasets.

In summary, heat stress elicited a highly conserved transcriptional programme characterised by suppression of biosynthetic and energetic functions. Although heatstroke shared the core molecular response to heat stress, it exhibited a distinct cerebellar and neuronal signature, suggesting engagement of pathogenic processes beyond a simple amplification of heat-shock pathways. Given that heatstroke is accompanied by systemic inflammation, circulatory dysfunction, coagulopathy, endothelial injury, and multi-organ stress,^11,43–45^ features that are largely absent from experimental heat-exposure models, the observed regional and cellular specificity may reflect the convergence of these additional injury mechanisms on a conserved heat-stress response.

As extreme heat events become increasingly frequent, defining the molecular basis of selective CNS vulnerability to heat-related injury is increasingly important. Heatstroke can occur in otherwise healthy individuals, in exertion and without exertion, and in people with existing neurological disorders that may be acutely and permanently aggravated by heat. It is an important cause of death during heatwaves, highlighting the need for further study in the context of ongoing climate change.

